# Testing the Toxicofera: comparative reptile transcriptomics casts doubt on the single, early evolution of the reptile venom system

**DOI:** 10.1101/006031

**Authors:** Adam D Hargreaves, Martin T Swain, Darren W Logan, John F Mulley

## Abstract

**Background:** The identification of apparently conserved gene complements in the venom and salivary glands of a diverse set of reptiles led to the development of the Toxicofera hypothesis – the idea that there was a single, early evolution of the venom system in reptiles. However, this hypothesis is based largely on relatively small scale EST-based studies of only venom or salivary glands and toxic effects have been assigned to only some of these putative Toxcoferan toxins in some species. We set out to investigate the distribution of these putative venom toxin transcripts in order to investigate to what extent conservation of gene complements may reflect a bias in previous sampling efforts.

**Results:** We have carried out the first large-scale test of the Toxicofera hypothesis and found it lacking in a number of regards. Our quantitative transcriptomic analyses of venom and salivary glands and other body tissues in five species of reptile, together with the use of available RNA-Seq datasets for additional species shows that the majority of genes used to support the establishment and expansion of the Toxicofera are in fact expressed in multiple body tissues and most likely represent general maintenance or “housekeeping” genes. The apparent conservation of gene complements across the Toxicofera therefore reflects an artefact of incomplete tissue sampling. In other cases, the identification of a non-toxic paralog of a gene encoding a true venom toxin has led to confusion about the phylogenetic distribution of that venom component.

**Conclusions:** Venom has evolved multiple times in reptiles. In addition, the misunderstanding regarding what constitutes a toxic venom component, together with the misidentification of genes and the classification of identical or near-identical sequences as distinct genes has led to an overestimation of the complexity of reptile venoms in general, and snake venom in particular, with implications for our understanding of (and development of treatments to counter) the molecules responsible for the physiological consequences of snakebite.

## Background

Snake venom is frequently cited as being highly complex or diverse [1–3] and a large number of venom toxin genes and gene families have been identified, predominantly from EST-based studies of gene expression during the re-synthesis of venom in the venom glands following manually-induced emptying (“milking”) [4–8] and some proteomic studies of extracted venom. It has been suggested that many of these gene families have originated via the duplication of a gene encoding a non-venom protein expressed elsewhere in the body followed by recruitment into the venom gland where natural selection can act to increase toxicity, with subsequent additional duplications leading to a diversification within gene families, often in a species-specific manner [9–11]. However, since whole genome duplication is a rare event in reptiles [12], the hypothesis that novelty in venom originates via the duplication of a “body” gene with subsequent recruitment into the venom gland requires both that gene duplication is a frequent event in the germline of venomous snakes and that the promoter and enhancer sequences that regulate venom gland-specific expression are relatively simple and easy to evolve. It also suggests a high incidence of neofunctionalisation rather than the more common process of subfunctionalisation [13–16].

The apparent widespread distribution of genes known to encode venom toxins in snakes in the salivary glands of a diverse set of reptiles, including both those that had previously been suggested to have secondarily lost venom in favour of constriction or other predation techniques and those that had previously been considered to have never been venomous led to the development of the Toxicofera hypothesis – the single, early evolution of venom in reptiles [17–19] (Figure 1). Analysis of a wide range of reptiles, including charismatic megafauna such as the Komodo dragon, *Varanus komodoensis* [20], has shown that the basal Toxicoferan venom system comprises at least 16 genes, with additional gene families subsequently recruited in different lineages [18, 19, 21].

**Figure 1.** Relationships of key vertebrate lineages and the placement of species discussed in this paper. A monophyletic clade of reptiles (which includes birds) is shaded green and the Toxicofera [21] are shaded red. Modified taxon names have been used for simplicity.

Although toxic effects have been putatively assigned to some Toxicoferan venom proteins in some species, the problem remains that their identification as venom components is based largely on their expression in the venom gland during venom synthesis and their apparent relatedness to other, known toxins in phylogenetic trees. It has long been known that all tissues express a basic set of “housekeeping” or maintenance genes [22] and it is therefore not surprising that similar genes might be found to be expressed in similar tissues in different species of reptiles and that these genes might group together in phylogenetic trees. However, the identification of transcripts encoding putative venom toxins in other body tissues would cast doubt on the classification of these Toxicoferan toxins as venom components, as it is unlikely that the same gene could fulfil toxic and non-toxic roles without evidence for alternative splicing to produce a toxic variant (as has been suggested for *acetylcholinesterase* in the banded krait, *Bungarus fasciatus* [11, 23]) or increased expression levels in the venom gland (where toxicity might be dosage dependent). In order to address some of these issues and to test the robustness of the Toxicofera hypothesis we have carried out a comparative transcriptomic survey of the venom or salivary glands, skin and cloacal scent glands of five species of reptile. Unlike the pancreas and other parts of the digestive system [24, 25], these latter tissues (which include a secretory glandular tissue (the scent gland) and a relatively inert, non-secretory tissue (skin)) have not previously been suggested to be the source of duplicated venom toxin genes and we would therefore only expect to find ubiquitous maintenance or “housekeeping” genes to be commonly expressed across these tissues. Study species included the venomous painted saw-scaled viper (*Echis coloratus*); the nonvenomous corn snake (*Pantherophis guttatus*) and rough green snake (*Opheodrys aestivus*) and a member of one of the more basal extant snake lineages, the royal python (*Python regius*). As members of the Toxicofera *sensu* Fry et al. [21] we would expect to find the basic Toxicoferan venom genes expressed in the venom or salivary glands of all of these species. In addition we generated corresponding data for the leopard gecko (*Eublepharis macularius*), a member of one of the most basal lineages of squamate reptiles that lies outside of the proposed Toxicofera clade (Figure 1). We have also taken advantage of available transcriptomes or RNA-Seq data for corn snake vomeronasal organ [26] and brain [27], garter snake (*Thamnophis elegans*) liver [28] and pooled tissues (brain, gonads, heart, kidney, liver, spleen and blood of males and females [29]), eastern diamondback rattlesnake (*Crotalus adamanteus*) and eastern coral snake (*Micrurus fulvius*) venom glands [7, 8, 30], king cobra (*Ophiophagus hannah*) venom gland, accessory gland and pooled tissues (heart, lung, spleen, brain, testes, gall bladder, pancreas, small intestine, kidney, liver, eye, tongue and stomach) [31], Burmese python (*Python molurus*) pooled liver and heart [32], green anole (*Anolis carolinensis*) pooled tissue (liver, tongue, gallbladder, spleen, heart, kidney and lung), testis and ovary [33] and bearded dragon (*Pogona vitticeps*), Nile crocodile (*Crocodylus niloticus*) and chicken (*Gallus gallus*) brains [27], as well as whole genome sequences for the Burmese python and king cobra [31, 34].

Assembled transcriptomes were searched for genes previously suggested to be venom toxins in *Echis coloratus* and related species [5, 35, 36] as well as those that have been used to support the Toxicofera hypothesis, namely *acetylcholinesterase, AVIT peptide* [9, 11, 18, 19, 23]*, complement c3/cobra venom factor, epididymal secretory protein* [19, 37], *c-type lectins* [38, 39]*, cysteine-rich secretory protein (crisp)* [40, 41]*, crotamine* [42, 43]*, cystatin* [44, 45]*, dipeptidylpeptidase, lysosomal acid lipase, renin aspartate protease* [5, 19, 35, 46]*, hyaluronidase* [47, 48]*, kallikrein* [49, 50]*, kunitz* [51]*, l-amino-acid oxidase* [52, 53]*, nerve growth factor* [54, 55]*, phospholipase A_2_* [15]*, phospholipase b* [7, 56, 57]*, ribonuclease* [58]*, serine protease* [59, 60]*, snake venom metalloproteinase* [61, 62]*, vascular endothelial growth factor* (*vegf*) [9, 17, 63, 64]*, veficolin* [65]*, vespryn, waprin* [19, 66–68] and *3-finger toxins* [69].

We find that many genes previously claimed to be venom toxins are in fact expressed in multiple tissues (Figure 2) and that transcripts encoding these genes show no evidence of consistently elevated expression level in venom or salivary glands compared to other tissues (Supplemental tables S5-S9). Only two putative venom toxin genes (*l-amino acid oxidase b2* and *PLA_2_ IIA-c*) showed evidence of a venom gland-specific splice variant across our multiple tissue data sets. We have also identified several cases of mistaken identity, where non-orthologous genes have been used to claim conserved, ancestral expression and instances of identical sequences being annotated as two distinct genes (see later sections). We propose that the putative ancestral Toxicoferan venom toxin genes do not encode toxic venom components in the majority of species and that the apparent venom gland-specificity of these genes is a side-effect of incomplete tissue sampling. Our analyses show that neither increased expression in the venom gland nor the production of venom-specific splice variants can be used to support continued claims for the toxicity of these genes.

**Figure 2.** Tissue distribution of proposed venom toxin transcripts. The majority of transcripts proposed to encode Toxicoferan venom proteins are expressed in multiple body tissues. Transcripts found in the assembled transcriptomes but which are assigned transcript abundance of <1 FPKM are shaded orange. Eco, painted saw-scaled viper (*Echis coloratus*); Pgu, corn snake (*Pantherophis guttatus*); Oae, rough green snake (*Opheodrys aestivus*); Pre, royal python (*Python regius*); Ema, leopard gecko (*Eublepharis macularius*). VG, venom gland; SAL, salivary gland; SCG, scent gland; SK, skin.

## Results

Based on our quantitative analysis of their expression pattern across multiple species, we identify the following genes as unlikely to represent toxic venom components in the Toxicofera. The identification of these genes as non-venom is more parsimonious than alternative explanations such as the reverse recruitment of a “venom” gene back to a “body” gene [70], which requires a far greater number of steps (duplication, recruitment, selection for increased toxicity, reverse recruitment) to have occurred in each species, whereas a “body” protein remaining a “body” protein is a zero-step process regardless of the number of species involved. The process of reverse recruitment must also be considered doubtful given the rarity of gene duplication in vertebrates (estimated to be between 1 gene per 100 to 1 gene per 1000 per million years [71–73].

### Acetylcholinesterase

We find identical *acetylcholinesterase* (*ache*) transcripts in the *E. coloratus* venom gland and scent gland (which we call transcript 1) and an additional splice variant expressed in skin and scent gland (transcript 2). Whilst the previously known splice variants in banded krait (*Bungarus fasciatus*) are differentiated by the inclusion of an alternative exon, analysis of the *E. coloratus ache* genomic sequence (accession number KF114031) reveals that the shorter transcript 2 instead comprises only the first exon of the *ache* gene, with a TAA stop codon that overlaps the 5’ GT dinucleotide splice site in intron 1. *ache* transcript 1 is expressed at a low level in the venom gland (6.60 FPKM) and is found in multiple tissues in all study species (Figure 2), as well as corn snake vomeronasal organ and garter snake liver. The shorter transcript 2 is found most often in skin and scent glands (Figure 2, Supplementary figure S1). The low expression level and diverse tissue distribution of transcripts of this gene suggest that *acetylcholinesterase* does not represent a Toxicoferan venom toxin. It should also be noted that the most frequently cited sources for the generation of a toxic version of *ache* in banded krait via alternative splicing include statements that *ache* “does not appear to contribute to the toxicity of the venom” [74], is “not toxic to mice, even at very high doses” [75] and is “neither toxic by itself nor acting in a synergistic manner with the toxic components of venom” [76].

### AVIT

We find only a single transcript encoding an AVIT peptide in our dataset, in the scent gland of the rough green snake (data not shown). The absence of this gene in all of our venom and salivary gland datasets, as well as the venom glands of the king cobra, eastern coral snake and Eastern diamondback rattlesnake and the limited number of sequences available on Genbank (one species of snake, *Dendroaspis polylepis* (accession number P25687) and two species of lizard, *Varanus varius* and *Varanus komodoensis* (accession numbers AAZ75583 and ABY89668 respectively)) despite extensive sampling, would suggest that it is unlikely to represent a conserved Toxicoferan venom toxin.

### Complement C3 (“cobra venom factor”)

We find identical transcripts encoding *complement c3* in all tissues in all species, with the exception of royal python skin (Figures 2 and 3) and we find only a single *complement c3* gene in the *E. coloratus* genome (data not shown). These findings, together with the identification of transcripts encoding this gene in the liver, brain, vomeronasal organ and tissue pools of various other reptile species (Figure 3) demonstrate that this gene does not represent a Toxicoferan venom toxin. However, the grouping of additional *complement c3* genes in the king cobra (*Ophiophagus hannah*) and monocled cobra (*Naja kaouthia*) in our phylogenetic tree does support a duplication of this gene somewhere in the Elapid lineage. One of these paralogs may therefore represent a venom toxin in at least some of these more derived species and the slightly elevated expression level of this gene in the venom or salivary gland of some of our study species suggests that *complement c3* has been exapted [77] to become a venom toxin in the Elapids. It seems likely that the identification of the non-toxic paralogue in other species (including veiled chameleon (*Chamaeleo calyptratus*), spiny-tailed lizard (*Uromastyx aegyptia*) and Mitchell's water monitor (*Varanus mitchelli*)) has contributed to confusion about the distribution of this “Cobra venom factor” (which should more rightly be called *complement c3b*), to the point where genes in alligator (*Alligator sinensis*), turtles (*Pelodiscus sinensis*) and birds (*Columba livia*) are now being annotated as venom factors (accession numbers XP 006023407-8, XP 006114685, XP 005513793, Figure 3).

**Figure 3.** Maximum likelihood tree of *complement c3* (“*cobra venom factor*”) sequences. Whilst most sequences likely represent housekeeping or maintenance genes, a gene duplication event in the elapid lineage (marked with *) may have produced a venom-specific paralog. An additional duplication (marked with +) may have taken place in *Austrelaps superbus*, although both paralogs appear to be expressed in both liver and venom gland. Geographic separation in king cobras (*Ophiophagus hannah*) from Indonesia and China is reflected in observed sequence variation. Numbers above branches are Bootstrap values for 500 replicates. Tissue distribution of transcripts is indicated using the following abbreviations: VG, venom gland; SK, skin; SCG, scent gland, AG, accessory gland; VMNO, vomeronasal organ and those genes found to be expressed in one or more body tissues are shaded blue.

### Cystatin

We find two transcripts encoding cystatins expressed in the venom gland of *E. coloratus* corresponding to *cystatin-e/m* and *f* (Supplementary figures S2 and S3). *cystatin-e/m* was found to be expressed in all tissues from all species used in this study (Figure 2), as well as corn snake vomeronasal organ and brain and garter snake liver and pooled tissues. The transcript encoding *cystatin-f* (which has not previously been reported to be expressed in a snake venom gland) is also expressed in the scent gland of *E. coloratus* and in the majority of other tissues of our study species. We find no evidence for a monophyletic clade of Toxicoferan cystatin-derived venom toxins and would agree with Richards et al. [45] that low expression level and absence of *in vitro* toxicity represents a “strong case for snake venom cystatins as essential housekeeping or regulatory proteins, rather than specific prey-targeted toxins…” Indeed, it is unclear why cystatins should be considered to be conserved venom toxins, since even from its earliest discovery in the venom of the puff adder (*Bitis arietans*) there has been “…no evidence that it is connected to the toxicity of the venom” [44].

### Dipeptidyl peptidases

We find identical transcripts encoding *dipeptidyl peptidase 3* and *4* in all tissues in all species except the leopard gecko (Figures 2, 4a and 4b), and both of these have a low transcript abundance in the venom gland of *E. coloratus*. *dpp4* is expressed in garter snake liver and Anole testis and ovary and *dpp3* is also expressed in garter snake liver, king cobra pooled tissues and Bearded dragon brain (Figures 4a and b). It is therefore unlikely the either *dpp3* or *dpp4* represent venom toxins.

**Figure 4.** Maximum likelihood tree of *dipeptidylpeptidase 3* (*dpp3*) and *dipeptidylpeptidase 4* (*dpp4*) sequences. Transcripts encoding *dpp3* and *dpp4* are found in a wide variety of body tissues, and likely represent housekeeping genes. Numbers above branches are Bootstrap values for 500 replicates. Tissue distribution of transcripts is indicated using the following abbreviations: VG, venom gland; SK, skin; SCG, scent gland, AG, accessory gland; VMNO, vomeronasal organ and those genes found to be expressed in one or more body tissues are shaded blue.

### Epididymal secretory protein

We find one transcript encoding epididymal secretory protein (ESP) expressed in the venom gland of *Echis coloratus* corresponding to type E1. This transcript is also found to be expressed at similar levels in the scent gland and skin of this species and orthologous transcripts are expressed in all three tissues of all other species used in this study (Figure 2 and Supplementary figure S4a), suggesting that this is a ubiquitously expressed gene and not a venom component. Previously described epididymal secretory protein sequences from varanids [78] and the colubrid *Cylindrophis ruffus* [21] do not represent *esp-e1* and their true orthology is currently unclear. However, our analysis of these and related sequences suggests that they are likely part of a reptile-specific expansion of esp-like genes and that the *Varanus* and *Cylindrophis* sequences do not encode the same gene (Supplementary figure S4b). Therefore there is not, nor was there ever, any evidence that epididymal secretory protein sequences represent venom components in the Toxicofera.

### Ficolin (“veficolin”)

We find one transcript encoding *ficolin* in the *E. coloratus* venom gland and identical transcripts in both scent gland and skin (Figure 2, Supplementary figure S5) and orthologus transcripts in all corn snake and leopard gecko tissues, as well as rough green snake salivary and scent glands and royal python salivary gland. Paralogous genes expressed in multiple tissues were also found in corn snake and rough green snake (Supplementary figure S5). These findings, together with additional data from available transcriptomes of pooled garter snake body tissues and bearded dragon and chicken brains show that *Ficolin* does not represent a Toxicoferan venom component.

### Hyaluronidase

Hyaluronidase has been suggested to be a “venom spreading factor” to aid the dispersion of venom toxins throughout the body of envenomed prey, and as such it does not represent a venom toxin itself [79]. We do however find two hyaluronidase genes expressed in the venom gland of *E. coloratus*. The first appears to be venom gland specific (based on available data) and has two splice variants including a truncated variant similar to sequences previously characterised from *Echis carinatus sochureki* (accession number DQ840262) and *Echis pyramidum leakeyi* (accession number DQ840255) venom glands [48]. Although we cannot rule out hyaluronidase in playing an active (but non-toxic) role in *Echis* venom, it is worth commenting that hyaluronan has been suggested to have a role in wound healing and the protection of the oral mucosa in human saliva [80]. The expression of hyaluronidases involved in hyaluronan metabolism in venom and/or salivary glands is therefore perhaps unsurprising.

### Kallikrein

We find two Kallikrein-like sequences in *E. coloratus*, one of which is expressed in all three tissues in this species (at a low level in the venom gland) and a variety of other tissues in the other study species, and one of which is found only in scent gland and skin (Figure 2, Supplementary figure S6). These genes do not represent venom toxins in *E. coloratus* and appear to be most closely-related to a group of mammalian <u>Kallikrein</u> (KLK) genes containing *KLK1*, *11*, *14* and *15* and probably represent the outgroup to a mammalian-specific expansion of this gene family. The orthology of previously published Toxicoferan Kallikrein genes is currently unclear and the majority of these sequences can be found in our serine protease tree (see later section and Supplementary figure S19).

### Kunitz

We find a number of transcripts encoding Kunitz-type protease inhibitors in our tissue data, with the majority of these encoding *kunitz1* and *kunitz2* genes (Figure 2 and Supplementary figure S7). The tissue distribution of these transcripts, together with the phylogenetic position of lizard and venomous snake sequences does not support a monophyletic clade of venom gland-specific Kunitz-type genes in the Toxicofera. The presence of protease inhibitors in reptile venom and salivary glands should perhaps not be too surprising and it again seems likely that the involvement of Kunitz-type inhibitors in venom toxicity in some advanced snake lineages (in this case mamba (*Dendroaspis sp.*) dendrotoxins and krait (*Bungarus multicinctus*) bungarotoxins [81, 82]) has led to confusion when non-toxic orthologs have been identified in other species.

### Lysosomal acid lipase

We find two transcripts encoding Lysosomal acid lipase genes in the *E. coloratus* venom gland transcriptome, one of which (*lipa-a*) is also expressed in skin and scent gland in this species and all three tissues in our other study species. *lipa-a*, despite not being venom gland specific, is more highly expressed in the venom gland (3,337.33 FPKM) than in the scent gland (484.49 FPKM) and skin (22.79 FPKM) of *E. coloratus,* although there is no evidence of elevated expression in the salivary glands of our other study species. As this protein is involved in lysosomal lipid hydrolysis [83] and the venom gland is a highly active tissue, we suggest that this elevated expression is likely related to high cell turnover. Transcripts of *lipa-b* are found at a low level in the venom and scent glands of *E. coloratus* and the scent gland of royal python (Figure 2, Supplementary figure S8). Neither *lipa-a* or *lipa-b* therefore encode venom toxins.

### Natriuretic peptide

We find only a single natriuretic peptide-like sequence in our dataset, in the skin of the royal python. The absence of this gene from the rest of our study species suggests that it is not a highly conserved Toxicoferan toxin.

### Nerve growth factor

We find identical transcripts encoding *nerve growth factor* (*ngf*) in all three *E. coloratus* tissues. Transcripts encoding the orthologous gene are also found in the corn snake salivary gland and scent gland; rough green snake scent gland and skin; royal python skin and leopard gecko salivary gland, scent gland and skin (Figure 2 and Supplementary figure S9). *ngf* is expressed at a higher level in the venom gland (525.82 FPKM) than in the scent gland (0.18 FPKM) and skin (0.58 FPKM) of *E. coloratus*, but not at an elevated level in the salivary gland of other species, again hinting at the potential for exaptation of this gene. Based on these findings, together with the expression of this gene in garter snake pooled tissues, we suggest that *ngf* does not encode a Toxicoferan toxin. However, we do find evidence for the duplication of *ngf* in cobras (Supplementary figure S9) suggesting that it may represent a venom toxin in at least some advanced snakes [84]. As with *complement c3*, it seems likely that the identification of non-toxic orthologs in distantly-related species has led to the conclusion that *ngf* is a widely-distributed venom toxin and confused its true evolutionary history.

### Phospholipase A_2_ (PLA_2_ Group IIE)

We find transcripts encoding Group IIE PLA_2_ genes in the venom gland of *E. coloratus* and the salivary glands of all other species (Figure 2, Supplementary figure S10). Although this gene appears to be venom and salivary-gland-specific (based on available data), its presence in all species (including the non-Toxicoferan leopard gecko) suggests that it does not represent a toxic venom component.

### Phospholipase B

We find a single transcript encoding *phospholipase b* expressed in all three *E. coloratus* tissues (Figures 2 and 5) and transcripts encoding the orthologous gene are found in all other tissues from all study species with the exception of rough green snake salivary gland. We also find *plb* transcripts in corn snake vomeronasal organ, garter snake liver, Burmese python pooled tissues (liver and heart) and bearded dragon brain (Figure 5). The two transcripts in each of rough green snake and corn snake are likely alleles or the result of individual variation, and actually represent a single *phospholipase b* gene from each of these species. Transcript abundance analysis shows this gene to be expressed at a low level in all tissues from all study species. Based on the phylogenetic and tissue distribution of this gene it is unlikely to represent a Toxicoferan venom toxin.

**Figure 5.** Maximum likelihood tree of *phospholipase b* (*plb*) sequences. Transcripts encoding *plb* are found in a wide variety of body tissues, and likely represent housekeeping genes. Numbers above branches are Bootstrap values for 500 replicates. Tissue distribution of transcripts is indicated using the following abbreviations: VG, venom gland; SK, skin; SCG, scent gland, AG, accessory gland; VMNO, vomeronasal organ and those genes found to be expressed in one or more body tissues are shaded blue.

### Renin (“renin aspartate protease”)

We find a number of transcripts encoding renin-like genes in the *E. coloratus* venom gland (Figures 2 and 6), one of which (encoding the canonical *renin*) is also expressed in the scent gland and is orthologous to a previously described sequence from the venom gland of the ocellated carpet viper (*Echis ocellatus*, accession number CAJ55260). We also find that the recently-published *Boa constrictor renin aspartate protease* (*rap*) gene (accession number JX467165 [21]) is in fact a *cathepsin d* gene, transcripts of which are found in all three tissues in all five of our study species. We suggest that this misidentification may be due to a reliance on BLAST-based classification, most likely using a database restricted to squamate or serpent sequences. It is highly unlikely that either *renin* or *cathepsin d* (or indeed any renin-like aspartate proteases) constitute venom toxins in *E. coloratus* or *E. ocellatus*, nor do they represent basal Toxicoferan toxins.

**Figure 6.** Maximum likelihood tree of renin-like sequences. Renin-like genes are expressed in a diversity of body tissues. The recently published *Boa constrictor* “RAP-Boa-1” sequence is clearly a *cathepsin d* gene and is therefore not orthologous to the *Echis ocellatus* renin sequence as has been claimed [21]. Numbers above branches are Bootstrap values for 500 replicates. Tissue distribution of transcripts is indicated using the following abbreviations: VG, venom gland; SK, skin; SCG, scent gland and those genes found to be expressed in one or more body tissues are shaded blue.

### Ribonuclease

Ribonucleases have been suggested to have a role in the generation of free purines in snake venoms [58] and the presence of these genes in the salivary glands of two species of lizard (*Gerrhonotus infernalis* and *Celestus warreni*) and two colubrid snakes (*Liophis peocilogyrus* and *Psammophis mossambicus*) has been used to support the Toxicofera [78, 85]. We did not identify orthologous ribonuclease genes in any of our salivary or venom gland data, nor do we find them in venom gland transcriptomes from the Eastern diamondback rattlesnake, king cobra and eastern coral snake (although we have identified a wide variety of other ribonuclease genes). The absence of these genes in seven Toxicoferans, coupled with the fact that they were initially described from only 2 out of 11 species of snake [85] and 3 out of 18 species of lizard [78] would cast doubt on their status as conserved Toxicoferan toxins.

### Three finger toxins (3ftx)

We find 2 transcripts encoding three finger toxin (3ftx)-like genes expressed in the *E. coloratus* venom gland, one of which is expressed in all 3 tissues (*3ftx-a*) whilst the other is expressed in the venom and scent glands (*3ftx-b*). Orthologous transcripts of *3ftx-a* are found to be expressed in all three tissues of corn snake, rough green snake salivary gland and skin, and royal python salivary gland. An ortholog of *3ftx-b* is expressed in rough green snake scent gland. We also find a number of different putative *3ftx* genes in our other study species, often expressed in multiple tissues (Figure 2, Supplementary figure S11). Based on the phylogenetic and tissue distribution of both of these genes we suggest that they do not represent venom toxins in *E. coloratus*. As with other proposed Toxicoferan genes such as *complement c3* and *nerve growth factor*, it seems likely that *3ftx* genes are indeed venom components in some species, especially cobras and other elapids [31, 69], and that the identification of their non-venom orthologs in other species has led to much confusion regarding the phylogenetic distribution of these toxic variants.

### Vespryn

We do not find *vespryn* transcripts in any *E. coloratus* tissues, although this gene is present in the genome of this species (accession number KF114032). We do however find transcripts encoding this gene in the salivary and scent glands of the corn snake, and skin and scent glands of the rough green snake, royal python and leopard gecko (Figure 2, Supplementary figure S12). We suggest that the tissue distribution of this gene in these species casts doubt on its role as a venom component in the Toxicofera.

### Waprin

We find a number of “waprin”-like genes in our dataset, expressed in a diverse array of body tissues. Our phylogenetic analyses (Supplementary figure S13) show that previously characterised “waprin” genes [8, 66, 68, 86, 87] most likely represent *WAP four-disulfide core domain 2* (*wfdc2*) genes which have undergone a squamate-specific expansion and that there is no evidence for a venom gland-specific paralog. It is unlikely therefore that these genes represent a Toxicoferan venom toxin. Indeed, the inland taipan (*Oxyuranus microlepidotus*) “Omwaprin” has been shown to be “…non-toxic to Swiss albino mice at doses of up to 10 mg/kg when administered intraperitoneally” [68] and is more likely to have an antimicrobial function in the venom or salivary gland.

## Implications for venom composition and complexity in *Echis coloratus*

The following genes show either a venom gland-specific expression or an elevated expression level in this tissue, but not both and as such we suggest that whilst they *may* represent venom toxins in *E. coloratus*, further analysis is needed in order to confirm this.

### Vascular endothelial growth factor

We find four transcripts encoding vascular endothelial growth factor (VEGF) expressed in the venom gland of *E. coloratus*. These correspond to *vegf-a*, *vegf-b*, *vegf-c* and *vegf-f* and of these *vegf-a*, *b* and *c* are also expressed in the skin and scent gland of this species (Figure 2). Transcripts encoding orthologs of these genes are expressed in all three tissues of all other species used in this study (with the exception of the absence of *vegf-a* in corn snake skin). In accordance with previous studies [7] we find evidence of alternative splicing of *vegf-a* transcripts in all species although no variant appears to be tissue-specific. It is likely that a failure to properly recognise and classify alternatively spliced *vegf-a* transcripts (Aird et al. 2013) may have contributed to an overestimation of snake venom complexity. *vegf-d* was only found to be expressed in royal python salivary gland and scent gland and all three tissues from leopard gecko (Figure 2, Supplementary figure S14). The transcript encoding VEGF-F is found only in the venom gland of *E. coloratus* and, given the absence of any Elapid *vegf-f* sequences in public databases as well as absence of this transcript in the two species of colubrid in our study, we suggest that *vegf-f* is specific to vipers. Whilst *vegf-f* has a higher transcript abundance in *E. coloratus* venom gland (186.73 FPKM) than *vegf-a* (3.24 FPKM), *vegf-b* (1.28 FPKM) and *vegf-c* (1.54 FPKM), compared to other venom genes in this species (see next section) it has a considerably lower transcript abundance suggesting it represents at most a minor venom component in *E. coloratus*.

### L-amino acid oxidase

We find transcripts encoding two *l-amino acid oxidase (laao)* genes in *E. coloratus*, one of which (*laao-b*) has two splice variants (Figure 2, Supplementary figure S15). *laao-a* transcripts are found in all three *E. coloratus* and leopard gecko tissues. *laao-b* is venom gland-specific in *E. coloratus* (based on the available data) and transcripts of the orthologous gene are found in the scent glands of corn snake, rough green snake and royal python. The splice variant *laao-b2* may represent a venom toxin in *E. coloratus* based on its specific expression in the venom gland of this species and elevated expression level (628.84 FPKM).

### Crotamine

We find a single *crotamine*-like transcript in *E. coloratus*, in the venom gland (Figure 2). Related genes are found in a variety of tissues in other study species (including the scent gland of the rough green snake, the salivary gland and skin of the leopard gecko, and in all three corn snake tissues), although the short length of these sequences precludes a definitive statement of orthology. This gene may represent a toxic venom component in *E. coloratus* based on its tissue distribution, but due to its low transcript abundance (10.95 FPKM) it is likely to play a minor role, if any.

The following genes are found only in the venom gland of *E. coloratus* and clearly show an elevated expression level (Figure 7). Whilst we classify these genes as encoding venom toxins in this species (Table 1) it should be noted that none of these genes support the monophyly of Toxicoferan venom toxins.

**Figure 7.** Graph of transcript abundance values of proposed venom transcripts in the *Echis coloratus* venom gland. The majority of Toxicoferan transcripts are expressed at extremely low level, with the most highly expressed genes falling into only four gene families (C-type lectins, Group IIA phospholipase A_2_, serine proteases and snake venom metalloproteinases). FPKM = Fragments Per Kilobase of exon per Million fragments mapped.

**Table 1.** Predicted venom composition of the painted saw-scaled viper, *Echis coloratus*

| Gene family |  | Number of genes |
| --- | --- | --- |
| SVMP |  | 13 |
| C-type lectin |  | 8 |
| Serine protease |  | 6 |
| PLA2 |  | 3 |
| CRISP |  | 1 |
| L-amino acid oxidase |  | 1 |
| VEGF |  | 1 |
| Crotamine |  | 1 |
| Total | 8 | 34 |

### Cysteine-rich secretory proteins (CRISPs)

We find transcripts encoding two distinct CRISPs expressed in the *E. coloratus* venom gland, one of which is also found in skin and scent gland (Figure 2). Phylogenetic analysis of these genes (which we call *crisp-a* and *crisp-b*) reveals that they appear to have been created as a result of a gene duplication event earlier in the evolution of advanced snakes (Supplementary Figure S16). *crisp-a* transcripts are also found in all three corn snake tissues, as well as rough green snake skin and scent gland and royal python scent gland. *crisp-b* is also found in corn snake salivary gland (Figure 2 and Supplementary figure S16) and the phylogenetic and tissue distribution of this gene suggest that it does indeed represent a venom toxin, produced via duplication of an ancestral *crisp* gene that was expressed in multiple tissues, including the salivary gland. The elevated transcript abundance of *crisp-b* (3,520.07 FPKM) in the venom gland of *E. coloratus* further supports its role as a venom toxin in this species (Figure 7). The phylogenetic and tissue distribution and low transcript abundance of *crisp-a* (0.61 FPKM in *E. coloratus* venom gland) shows that it is unlikely to be a venom toxin. We also find no evidence of a monophyletic clade of reptile venom toxins and therefore suggest that, contrary to earlier reports [20, 78], the CRISP genes of varanid and helodermatid lizards do not represent shared Toxicoferan venom toxins and, if they are indeed toxic venom components, have been recruited independently from those of the advanced snakes. Regardless of their status as venom toxins, it appears likely that the diversity of CRISP genes in varanid lizards in particular [17] has been overestimated as a result of the use of negligible levels of sequence variation to classify transcripts as representing distinct gene products (Supplementary figures S23 and S24).

### C-type lectins

We find transcripts encoding 11 distinct C-type lectin genes in the *E. coloratus* venom gland, one of which (*ctl-a*) is also expressed in the scent gland of this species. The remaining 10 genes (*ctl-b* to *k*) are found only in the venom gland and form a clade with other viper C-type lectin genes (Figure 2, Supplementary figure S17). Of these, 6 are highly expressed in the venom gland (*ctl-b* to *d*, *ctl-f* to *g* and *ctl-j*) with a transcript abundance range of 3,706.21-24,122.41 FPKM (Figure 7). The remainder of these genes (*ctl-e*, *ctl-h* to *i* and *ctl-k*) show lower transcript abundance (0.80-1,475.88 FPKM), with two (*ctl-i* and *k*) being more lowly expressed than *ctl-a* (230.06 FPKM). A number of different C-type lectin genes are found in our other study species, often expressed in multiple tissues (Supplementary figure S17). We suggest therefore that the 6 venom-gland specific C-type lectin genes which are highly expressed are indeed venom toxins in *E. coloratus* and that these genes diversified via the duplication of an ancestral gene with a wide expression pattern, including in salivary/venom glands. Based on their selective expression in the venom gland (from available data) the remaining four C-type lectin genes cannot be ruled out as putative toxins, although their lower transcript abundance suggests that they are likely to be minor components in *E. coloratus* venom. It should also be noted that a recent analysis of king cobra (*Ophiohagus hannah*) venom gland transcriptome and proteome suggested that “…lectins do not contribute to king cobra envenoming” [31].

### Phospholipase A_2_ (PLA_2_ Group IIA)

We find five transcripts encoding Group IIA PLA_2_ genes in *E. coloratus*, three of which are found only in the venom gland and two of which are found only in the scent gland (these latter two likely represent intra-individual variation in the same transcript) (Figure 2, Supplementary figure S18). The venom gland-specific transcript *PLA_2_ IIA-c* is highly expressed (22,520.41 FPKM) and likely represents a venom toxin, and may also be a putative splice variant although further analysis is needed to confirm this. *PLA_2_ IIA-d* and *IIA-e* show an elevated, but lower, expression level (1,677.15 FPKM and 434.67 FPKM respectively, Figure 7). Based on tissue and phylogenetic distribution we would propose that these three genes may represent putative venom toxins (Table 1).

### Serine proteases

We find 6 transcripts encoding Serine proteases in *E. coloratus* (Figure 2, Supplementary figure S19) which (based on available data) are all venom gland specific. Four of these transcripts are highly expressed in the venom gland (*serine proteases a-c* and *e*; 3,076.01-7,687.03 FPKM) whilst two are expressed at a lower level (*serine proteases d* and *f*; 1,098.45 FPKM and 102.34 FPKM respectively, Figure 7). Based on these results we suggest *serine proteases a, b, c* and *e* represent venom toxins whilst *serine proteases d* and *f* may represent putative venom toxins (Table 1).

### Snake venom metalloproteinases

We find 21 transcripts encoding snake venom metalloproteinases in *E. coloratus* and of these 14 are venom gland-specific, whilst another (*svmp-n*) is expressed in the venom gland and scent gland. Five remaining genes are expressed in the scent gland only whilst another is expressed in the skin (Figure 2, Supplementary figure S20). Of the 14 venom gland-specific SVMPs we find 4 to be highly expressed (5,552.84-15,118.41 FPKM, Figure 7). In the absence of additional data, we classify the 13 venom gland-specific *svmp* genes as venom toxins in this species (Table 1).

## Discussion

Our transcriptomic analyses have revealed that all 16 of the basal venom toxin genes used to support the hypothesis of a single, early evolution of venom in reptiles (the Toxicofera hypothesis [17–21, 78]), as well as a number of other genes that have been proposed to encode venom toxins in multiple species are in fact expressed in multiple tissues, with no evidence for consistently higher expression in venom or salivary glands. Additionally, only two genes in our entire dataset of 74 genes in five species were found to encode possible venom gland-specific splice variants (*l-amino acid oxidase b2* and *PLA_2_ IIA-c*). We therefore suggest that many of the proposed basal Toxicoferan genes most likely represent housekeeping or maintenance genes and that the identification of these genes as conserved venom toxins is a side-effect of incomplete tissue sampling. This lack of support for the Toxicofera hypothesis therefore prompts a return to the previously held view [88] that venom in different lineages of reptiles has evolved independently, once at the base of the advanced snakes, once in the helodermatid (gila monster and beaded lizard) lineage and, possibly, one other time in monitor lizards, although evidence for a venom system in this latter group [20, 78, 89] may need to be reinvestigated in light of our findings. The process of reverse recruitment [70], where a venom gene undergoes additional gene duplication events and is subsequently recruited from the venom gland back into a body tissue (which was proposed on the basis of the placement of garter snake and Burmese python “physiological” genes within clades of “venom” genes) must also be re-evaluated in light of our findings.

Bites by venomous snakes are thought to be responsible for as many as 1,841,000 envenomings and 94,000 deaths annually, predominantly in the developing world [90, 91] and medical treatment of snakebite is reliant on the production of antivenoms containing antibodies, typically from sheep or horses, that will bind and neutralise toxic venom proteins [92]. Since these antivenoms are derived from the injection of crude venom into the host animal they are not targeted to the most pathogenic venom components and therefore also include antibodies to weakly- or non-pathogenic proteins, requiring the administration of large or multiple doses [11], increasing the risks of adverse reactions. A comprehensive understanding of snake venom composition is therefore vital for the development of the next generation of antivenoms [2, 11, 93] as it is important that research effort is not spread too thinly through the inclusion of non-toxic venom gland transcripts. Our results suggest that erroneous assumptions about the single origination and functional conservation of venom toxins across the Toxicofera has led to the complexity of snake venom being overestimated by previous authors, with the venom of the painted saw-scaled viper, *Echis coloratus* likely consisting of just 34 genes in 8 gene families (Table 1, based on venom gland-specific expression and a ‘high’ expression, as defined by presence in the top 25% of transcripts [94] in at least two of four venom gland samples), fewer than has been suggested for this and related species in previous EST or transcriptomic studies [5, 35]. However, it is noteworthy that the results of our analyses accord well with proteomic analyses of venom composition in snakes, which range from an almost identical complement of 35 toxins in 8 gene families for the related ocellated carpet viper, *Echis ocellatus* [36] to between 24-61 toxins in 6-14 families in a range of other species (Table 2). Far from being a “complex cocktail” [10, 11, 95, 96], snake venom may in fact represent a relatively simple mixture of toxic proteins honed by natural selection for rapid prey immobilisation, with limited lineage-specific expansion in one or a few particular gene families.

**Table 2.** Predicted numbers of venom toxins and venom toxin families from proteomic studies of snake venom accord well with our transcriptome results.

| Species | Number of toxins | Number of toxin families |
| --- | --- | --- |
| *Bitis caudalis* [103] | 30 | 8 |
| *Bitis gabonica gabonica* [104] | 35 | 12 |
| *Bitis gabonica rhinoceros* [103] | 33 | 11 |
| *Bitis nasicornis* [103] | 28 | 9 |
| *Bothriechis schlegelii* [105] | ? | 7 |
| *Cerastes cerastes* [106] | 25-30 | 6 |
| *Crotalus atrox* [107] | ~24 | ~9 |
| *Echis ocellatus* [36] | 35 | 8 |
| *Lachesis muta* [108] | 24-26 | 8 |
| *Naja kaouthia* [109] | 61 | 12 |
| *Ophiophagus hannah* [31] | ? | 14 |
| *Vipera ammodytes* [110] | 38 | 9 |

In order to avoid continued overestimation of venom complexity, we propose that future transcriptome-based analyses of venom composition must include quantitative comparisons of multiple body tissues from multiple individuals and robust phylogenetic analysis that includes known paralogous members of gene families. We would also encourage the use of clearly explained, justifiable criteria for classifying highly similar sequences as new paralogs rather than alleles or the result of PCR or sequencing errors, as it seems likely that some available sequences from previous studies have been presented as distinct genes on the basis of extremely minor (or even non-existent) sequence variation (see Supplementary figures S21-S24 for examples of identical or nearly identical ribonuclease and CRISP sequences and Supplementary figures S25 and S26 for examples of the same sequence being annotated as two different genes). As a result, the diversity of “venom” composition in these species may have been inadvertently inflated.

Additionally, we would encourage the adoption of a standard nomenclature for reptile genes, as the overly-complicated and confusing nomenclature used currently (Table 3) may also contribute to the perceived complexity of snake venom. We propose that such a nomenclature system should be based on the comprehensive standards developed for Anole lizards [97], for example:

- “Gene symbols for all…species should be written in lower case only and in italics, e.g., *gene2*.”
- “Whenever criteria for orthology have been met… the gene symbol should be comparable to the human gene symbol, e.g., if the human gene symbol is *GENE2*, then the gene symbol would be *gene2*.”
- “Duplication of the ortholog of a mammalian gene will be indicated by an “a” or “b” suffix, e.g., *gene2a* and *gene2b*. If the mammalian gene symbol already contains a suffix letter, then there would be a second letter added, e.g., *gene4aa* and *gene4ab*.”

It seems likely that the application of our approach to other species (together with proteomic studies of extracted venom) will lead to a commensurate reduction in claimed venom diversity, with clear implications for the development of next generation antivenoms: since most true venom genes are members of a relatively small number of gene families, it is likely that a similarly small number of antibodies may be able to bind to and neutralise the toxic venom components, especially with the application of “string of beads” techniques [98] utilising fusions of short oligopeptide epitopes designed to maximise the cross-reactivity of the resulting antibodies [2].

**Table 3.**
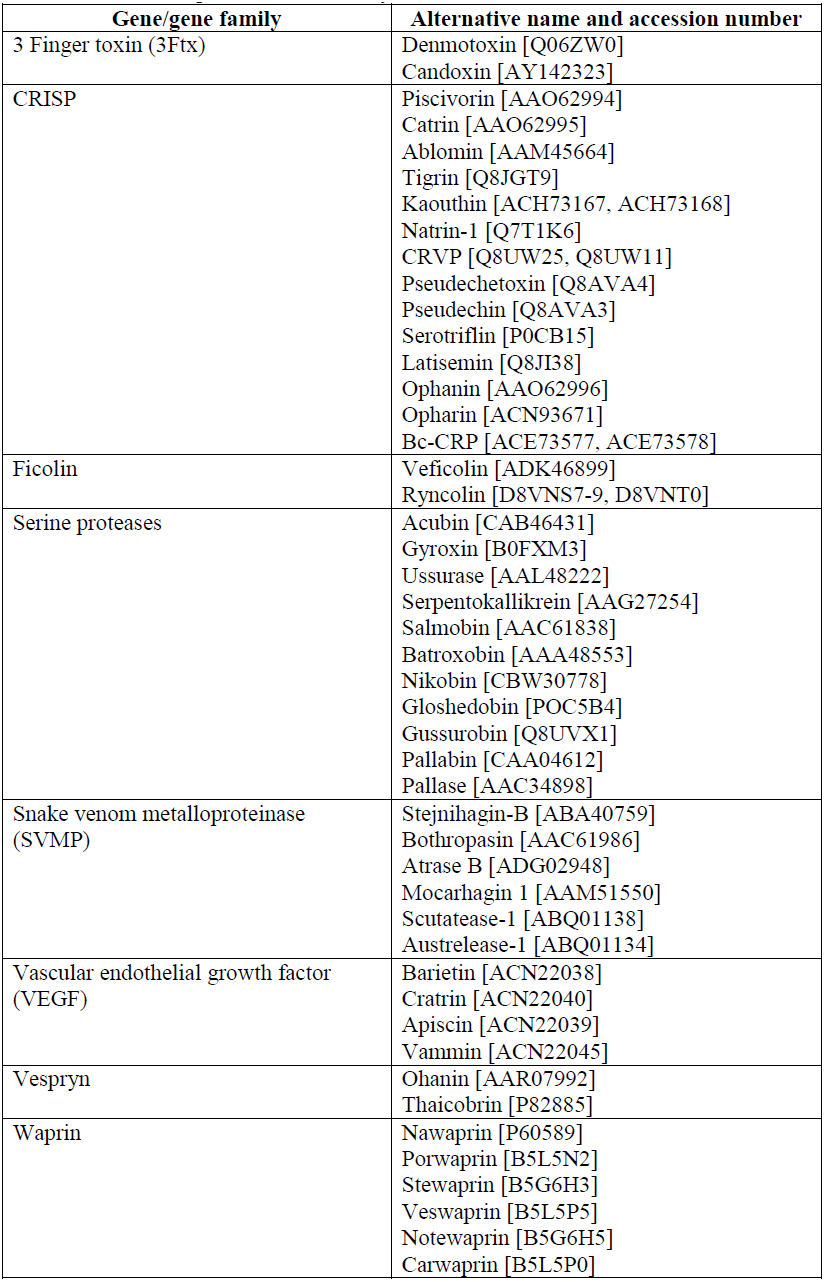
Venom gene nomenclature. Lack of a formal set of nomenclatural rules for venom toxins has led to an explosion of different gene names and may have contributed to the overestimation of reptile venom diversity.

## Conclusions

We suggest that identification of the apparently conserved Toxicofera venom toxins in previous studies is most likely a side effect of incomplete tissue sampling, compounded by incorrect interpretation of phylogenetic trees and the use of BLAST-based gene identification methods. It should perhaps not be too surprising that homologous tissues in related species would show similar gene complements and the restriction of most previous studies to only the “venom” glands means that monophyletic clades of reptile sequences in phylogenetic trees have been taken to represent monophyletic clades of venom toxin genes. Whilst it is true that some of these genes do encode toxic proteins in some species (indeed, this was often the basis for their initial discovery) the discovery of orthologous genes in other species does not necessarily demonstrate shared toxicity. In short, toxicity in one does not equal toxicity in all.

## Methods

Experimental methods involving animals followed institutional and national guidelines and were approved by the Bangor Universty Ethical Review Committee.

### RNA-Seq

Total RNA was extracted from four venom glands taken from four individual specimens of adult Saw-scaled vipers (*Echis coloratus*) at different time points following venom extraction in order to capture the full diversity of venom genes (16, 24 and 48 hours post-milking). Additionally, total RNA from two scent glands and two skin samples of this species and the salivary, scent glands and skin of two adult corn snakes (*Pantherophis guttatus*), rough green snakes (*Opheodrys aestivus*), royal pythons (*Python regius*) and leopard geckos (*Eublepharis macularius*) was also extracted using the RNeasy mini kit (Qiagen) with on-column DNase digestion. Only a single corn snake skin sample provided RNA of high enough quality for sequencing. mRNA was prepared for sequencing using the TruSeq RNA sample preparation kit (Illumina) with a selected fragment size of 200-500bp and sequenced using 100bp paired-end reads on the Illumina HiSeq2000 or HiSeq2500 platform.

### Quality control, assembly and analysis

The quality of all raw sequence data was assessed using FastQC [99] and reads for each tissue and species were pooled and assembled using Trinity [100] (sequence and assembly metrics are provided in Supplemental tables S1-S3). Putative venom toxin amino acid sequences were aligned using ClustalW [101] and maximum likelihood trees constructed using the Jones-Taylor-Thornton (JTT) model with 500 Bootstrap replicates. Transcript abundance estimation was carried out using RSEM [102] as a downstream analysis of Trinity (version trinityrnaseq_r2012-04-27). Sets of reads were mapped to species-specific reference transcriptome assemblies (Supplementary table S4) to allow comparison between tissues on a per-species basis and all results values shown are in FPKM (<u>F</u>ragments <u>P</u>er <u>K</u>ilobase of exon per <u>M</u>illion fragments mapped). Individual and mean FPKM values for each gene per tissue per species are given in Supplementary tables S5-S9. All transcript abundance values given within the text are based on the average transcript abundance per tissue per species to account for variation between individual samples.

Transcriptome reads were deposited in the European Nucleotide Archive (ENA) database under accession #ERP001222 and GenBank under accession numbers XXX and XXX and genes used to reconstruct phylogenies are deposited in GenBank under accession numbers XXXX-XXXX

## Authors' contributions

JFM, ADH and DWL designed the experiments and all authors carried out them out. JFM and ADH prepared the manuscript and this was seen, modified and approved by all authors.

## Acknowledgements

The authors wish to thank A. Tweedale, R. Morgan, G. Cooke, A. Barlow, C. Wüster and M. Hegarty for technical assistance and N. Casewell, W. Wüster and A. Malhotra for useful discussions. We are grateful to the staff of High Performance Computing (HPC) Wales for enabling and supporting our access to their systems. This research was supported by a Royal Society Research Grant awarded to JFM (grant number RG100514) and Wellcome trust funding to DWL (grant number 098051). JFM and MTS are supported by the Biosciences, Environment and Agriculture Alliance (BEAA) between Bangor University and Aberystwyth University and ADH is funded by a Bangor University 125^th^ Anniversary Studentship.

